# Molecular mechanism of acid stress response of *A. acidoterrestris* DSM 3922^T^ under sublethal pH environment

**DOI:** 10.1101/2023.07.13.548807

**Authors:** Xiaoxue Liu, Youzhi Wu, Lingxia Jiao, Junjian Ran, Linjun Sun, Fuzhou Ye, Xinhong Liang, Ruixiang Zhao

## Abstract

Acid-responsive proteome expression profiles of *Alicyclobacillus acidoterrestris* (*A. acidoterrestris*) were analysed using label-free quantitative mass spectrometry to investigate its acid resistance mechanism at sublethal pH. Totally, 325 differential expression proteins were identified during acid stress at pH2.5 condition for 15 min, of which the expressions of 83 proteins were up-regulated and the other 242 proteins expressions were down-regulated. Differentially expressed proteins were mainly involved in organic nitrogen compounds metabolism, small molecule metabolism, organic acid metabolism and signal transduction. Overall, they were mapped into 97 metabolic pathways. Combination of KEGG pathway analysis and protein functional analysis suggested that the pH homeostasis system, changes in metabolic pathways, cell membrane permeability and DNA repair are the main acid resistance mechanisms of *A. acidoterrestris* at sublethal pH conditions. It is speculated that *A. acidoterrestris* may sense and transmit pH signals from the external environment through the nhaB protein which holds histidine-dependent acid resistance system, initiating a series of acid tolerance reactions. Our study demonstrated global physiological response of *A. acidoterrestris* to sublethal pH, which provided a better understanding of acid adaption mechanism of *A. acidoterrestris*.

## Introduction

*A. acidoterrestris* are capable of growing at pH range from 3.0 to 6.0 (Yokota, ujii, & Goto, 2007) and could cause spoilage of acidic beverages due to its acidophilic and thermophilic resistance (Smit, Cameron, & Venter, 2011). Since *A. acidoterrestris* was isolated in 1984 (Cerny, Hennlich, & Poralla, 1984), spoilage incidents caused by *A. acidoterrestris* have increased and more diverse types of products including orange juice, tomato juice, pineapple juice, grape juice, canned tomatoes and iced tea etc. have become contaminated (Alpas, Alma, & Bozoglu, 2003; McKnight, Eiroa, Sant’Ana, & Massaguer, 2010; Oteiza, Soto, Alvarenga, Sant’Ana, & Giannuzzi, 2014; Walker & Phillips, 2005; Yue, Zhang, & Yuan, 2014). It has caused huge economic losses to fruit juice industry and become an industry-wide problem (Cai et al., 2015; Chang & Kang, 2004; Kapetanakou, Passiou, & Chalkou, 2020). The studies indicated that elimination of contamination by *A. acidoterrestris* was key to preventing acidic beverages spoilage. International trade required that *A. acidoterrestris* in apple condensed juice should not more than 1/10 (Chang & Kang, 2004). Therefore, the control of *A. acidoterrestris* hazard has become a common concern in the field of food research and fruit juice industry. Many research was focused on the detection and control of *A. acidoterrestris* in fruits juice (Mast, Dietrich, Didier, & Martlbauer, 2016; A. M. Ribeiro, Vanetti, & Paiva, 2022; L. R. Ribeiro & Cristianini, 2021).

Although the heat and acid resistance mechanism of *A. acidoterrestris* has been explored from the aspects of chemical composition of cell membrane (Moore, Walker, & Tornus, 1997; Wisotzkey, Jurtshuk, & Fox, 1992), thermal stability of DNA (Shemesh, Pasvolsky, Sela, Green, & Zakin, 2013; Wisotzkey et al., 1992) and functional characteristics of other macromolecules, especially some enzymes (Zhao, Jiao, & Xu, 2022; Zhao et al., 2021) and heat shock proteins (L. Jiao, Fan, Hua, Wang, & Wei, 2012; L. X. Jiao, Ran, Xu, & Wang, 2015; X. Xu et al., 2017), the perceptual and regulatory mechanism of *A. acidoterrestris* to high temperature and low pH remains poorly understood. As an obligate acidophilic bacterium, *A. acidoterrestris* might have evolved its unique acid tolerance mechanism. Proteomics studies can analyse the dynamic changes of bacterial intracellular proteins in response to the environment stress and have become a new strategy to explore the mechanism of bacterial stress response. Our work on the proteomics changes of *A. acidoterrestris* under acid stress reveal the key regulator protein involved in stress response and elucidate the regulation mechanism of *A. acidoterrestris* in responding to acid stress. This study will contribute to fully reveal the acid tolerance mechanism of *A. acidoterrestris* from the protein level and provide a theoretical and experimental basis for acidic beverages preservation as well as microbial acid-resistance research.

## Materials and methods

### Strain, media and culture conditions Bacterial strain

*Alicyclobacillus acidoterrestris* DSM 3922^T^ was purchased from Leibniz Institute DSMZ (German Collection of Microorganisms and Cell Cultures, Braunschweig, Germany). It was confirmed that the strain was *A. acidoterrestris* by 16S rRNA sequence analysis.

### Culture medium

AAM (*A. acidoterrestris* medium) medium was prepared as described by L. X. Jiao et al. (2015). In this study, AAM medium (different pH as stated) was used as the culture medium for *A. acidoterrestris* DSM 3922^T^.

### Growth and acid stress conditions

The strain of *A. acidoterrestris* DSM 3922^T^ was activated in AAM medium for 16 h at 45 °C. Then 0.5 ml cell suspension [final concentration 1% (v/v) as the inoculum] was added into 50 mL fresh AAM (pH4.0) medium in a 250 mL conical flask and incubated in a thermostatic shaking water bath (250 r/min) at 45 °C until reach the logarithmic phase. Five millilitres of the bacterial cultures were used as control and the other were acid-shocked by resuspending bacteria in AAM medium with pH3.0, 2.5, 2.0, respectively. Each of those different pH samples together with control group, culture under same condition for 15-, 30-, 45-, and 60-min duration correspondingly.

### Detection of bacterial viability

After acid stress, the bacterial viability and its morphology of *A. acidoterrestris* were examined by plate counting method and scanning electron microscope (SEM), respectively. The plate counting method was based on the method described by X. Xu et al. (2017), the acid-treated samples of 1 mL were taken and serial dilutions of the treated samples were prepared in a stroke-physiological saline solution, then 100 μL of serial dilutions samples plated in triplicate onto AAM plates. Plates were incubated for 24 h at 45 °C and colony forming units were enumerated to estimate cell viability. For SEM to observe the morphology of *A. acidoterrestris*, according to the method described by Hsueh et al. (2012), bacteria samples that were washed five times by ultrapure sterile water were fixed with 2.5% glutaraldehyde, dehydrated with graded concentrations of ethanol, air dried and gold coated. The samples were then examined by SEM.

### Sample preparation for proteomics analysis

After acid stress, bacteria samples of control and experimental groups (acid shock) were collected by centrifugation at 5000×g for 10 min. The samples were washed twice with phosphate-buffered saline (PBS), pH 7.0, and then suspended in lysis buffer (SDT) containing 4% (w/v) SDS, 1 mM DTT, 100 mM Tris-HCl pH7.6. After grind and ultrasonic lysis, the supernatants were collected by centrifugation and then protein quantification was estimated using BCA method.

Filter-aided sample preparation (FASP) method was used to convert proteins into peptides as described by Wiśniewski and colleague (Wisniewski, Zougman, Nagaraj, & Mann, 2009). For each sample, totally 200 μg of protein was proteolyzed. Briefly, DTT was added to 200 μg protein to the final concentration of 100 mM, incubated at 56 °C for 1 h and cooled down to room temperature. After that, 200 μL UA buffer (8 M Urea, 150 mM Tris-HCl pH 8.5) and peptide solution were transferred to 10 kDa filter for removing other low-molecular-weight components. Finally, the protein suspension was digested with 4 μg trypsin (Promega) in 40 μL NH_4_HCO_3_ (50 mM) at 37 °C for 16h. The resulting peptides were collected by centrifugation. The peptide content was determined using BCA method (Pierce™ Quantitative Colorimetric Peptide Assay).

### Peptide samples analyses by NanoLC-MS/MS

The nano-RPLC (reverse phase-high performance liquid chromatography, RP-HPLC) separation was performed on EASY-nLC1000 system (Thermo Fisher Scientific). Mobile phase A and B were 0.1% formic acid in water and 0.1% formic acid in acetonitrile, respectively. Briefly, 1 μg of peptide samples were loaded onto a C18 reversed□phase trap column (100 μm × 2 cm, Thermo Fisher Scientific) equilibrated with 100% A phase. Then, samples were eluted from the C18 analytical column (75 μm × 15 cm, Thermo Fisher Scientific) at a flow rate of 300 nL/min with the linear gradient increase from 5 to 40% B (130 min). At 140 min, the gradient increased to 90% B and was held there for 10 min. At 160 min, the gradient returned to 5% B to re-equilibrate the column for the next injection.

Eluting peptides were directly analysed via tandem mass spectrometry (MS/MS) on an LTQ-OrbitrapVelos Pro mass spectrometer (Thermo Finnigan, San Jose, CA) equipped with a nanoelectrospray ion source and MS data were acquired in data-dependent mode. A spray voltage of 1.8 kV and an ion transfer tube temperature of 250°C were applied. The instrument was calibrated using standard compounds and the MS spectra were acquired in a data-dependent manner in the scanning range of 350-1800 m/z. The MS/MS data were acquired in the linear ion trap by targeting top 10 most abundant ions for fragmentation using low-energy collision-induced dissociation experiments (using a normalized collision energy of 35%, an activation q of 0.25, and an activation time of 30 ms). Dynamic exclusion was set to a repeat count of 1 with a 30 s duration. The resolution for full MS scan analyses was 60, 000 at m/z 400. MS scans were recorded in profile mode and the MS/MS was recorded in centroid mode to reduce data file size.

### MS data analysis and evaluation

The raw data files acquired from mass spectrometry runs were analyzed using MaxQuant software (version 1.3.0.5, Max-Planck-Institute of Biochemistry, Am Klopferspitz, Germany). Enzyme specification during the search was trypsin. Carbamidomethylation (CAM) of cysteine was selected as a fixed modification, while oxidation of methionine and N-terminal acetylation were selected as variable modification. The search parameters used were as follows: 20 ppm tolerance for precursor ion mass, 0.5 Da for fragment ion mass and 6 ppm for the main search, respectively. Tandem MS search was done using the Andromeda search engine integrated into Maxquant and was run against target databases against the UniProt database (uniprot_*Alicyclobacillus acidoterrestris*_4110 _20160929.fasta). Minimum cutoff for peptide length was set at 7 amino acids and two missed cleavages were permitted.

The peptide and protein FDR (false discovery rate) was controlled at 0.01. A minimum of one sequence-unique peptides was required for identification. Validated peptides were grouped into individual protein clusters by Maxquant software. Feature matching between runs was done with a retention time window of 2 min and the Label-free quantitation (LFQ) function was enabled. The Maxquant peptide and protein quantification results from the ‘peptides.txt’ and ‘proteinGroups.txt’ files were imported into Perseus software (version 1.5.1.6) for further analysis.

### Protein quantification

According to the method described by Luber et al. (2010), protein abundance was calculated based on the normalized spectral protein intensity (label-free quantitation intensity, LFQ intensity) and the relative protein ratio between the groups was calculated by comparing the average abundance values of the protein in each group. Abundance changes morn than 1.5-fold or less than 0.67-fold compared with control with a significant change (p<0.05) was used as the thresholds to define differentially expressed proteins according to the criteria reported by Chen, Luo, Ding, and Xu (2016) and Le Bihan, Rayner, Roy, and Spagnolo (2013).

### Bioinformatics analysis of proteomics data

The identified proteins were classified according to annotations from the UniProt knowledgebase (http://www.uniprot.org/). OmicsBean (http://www.omicsbean.cn), a multi-omics data analysis tool was used to conduct Gene Ontology (GO) analysis of differentially abundant proteins. Kyoto Encyclopedia of Genes and Genomes (KEGG) pathway analysis (http://www.genome.ad.jpkegg/pathway.html) was performed to enrich high-level functions in the defined biological systems.

### Quantitative RT-qPCR validation

The potential key genes related to acid stress were validated using real-time quantitative RT-qPCR. RNAfast1000 (Pioneer Biotechnology, Inc) and RevertAid™ First Strand cDNA Synthesis Kit (Fermentas, Lithuania) were used for total RNA isolation and reverse transcription. Real-time quantitative PCR was performed using a Maxima SYBR Green/ROX qPCR Master Mix (Thermo Fisher Scientific, USA) and the TL988 Real-time Quantitative PCR Detection System (Tianlong, Tianjin, China). The RT-qPCR data were analysed using the 2^− ΔΔCT^ method described by Livak and Schmittgen (Livak & Schmittgen, 2001). The following thermal protocol of PCR reaction was used: pre-denaturation at 95 °C for 4 min, initial denaturation at 95 °C for 30 s, amplification with denaturation at 95 °C for 5 s and annealing and extension at 60 °C for 31 s for 35 cycles in total. The qRT-PCR data were analyzed using the 2^−ΔΔCt^ method described by Livak and Schmittgen (2001)

### Data Processing and Statistical Analysis

Three biological and two technical replications of each sample were analyzed in order to assess quantitative reproducibility and missing data by label-free approaches. Statistical significance between the control and experimental group was evaluated using two pairs t-test analysis with a *P*-value cutoff of 0.05. The level of significance was set at a p-value of less than 0.05 or 0.01.

## Results

### Effect of acid stress on cell viability of *A. acidoterrestris*

When *A. acidoterrestris* was transferred from the optimum growth pH (pH4.0) to the minimum growth pH (pH3.0), bacteria continued to grow and reproduce themselves but with the reduction of bacterial numbers even 15 min duration under pH3.0 condition compared with the control group of pH4.0 (Fig. 1). Additionally, *A. acidoterrestris* that cultured at pH2.5 and pH2.0 condition, the number of viable bacteria showed a further decreasing trend (Fig. 1). Furthermore, the surface of *A. acidoterrestris* became uneven and the bacteria cells adhere to each other (Fig. 2), indicating that the bacteria phenotype change under acid stress. When *A. acidoterrestris* were treated at pH2.5 for 15 min, the bacteria quantity decreased slightly, and the cell damage was lighter compared with pH2.0 condition. It is speculated that the bacteria under acid stress of pH2.5 were in its survival state and the intracellular protein expression could have changed. So finally, the sample acid-shocked at pH2.5 for 15 min was selected to conduct the proteomics analysis.

**Fig 1.**
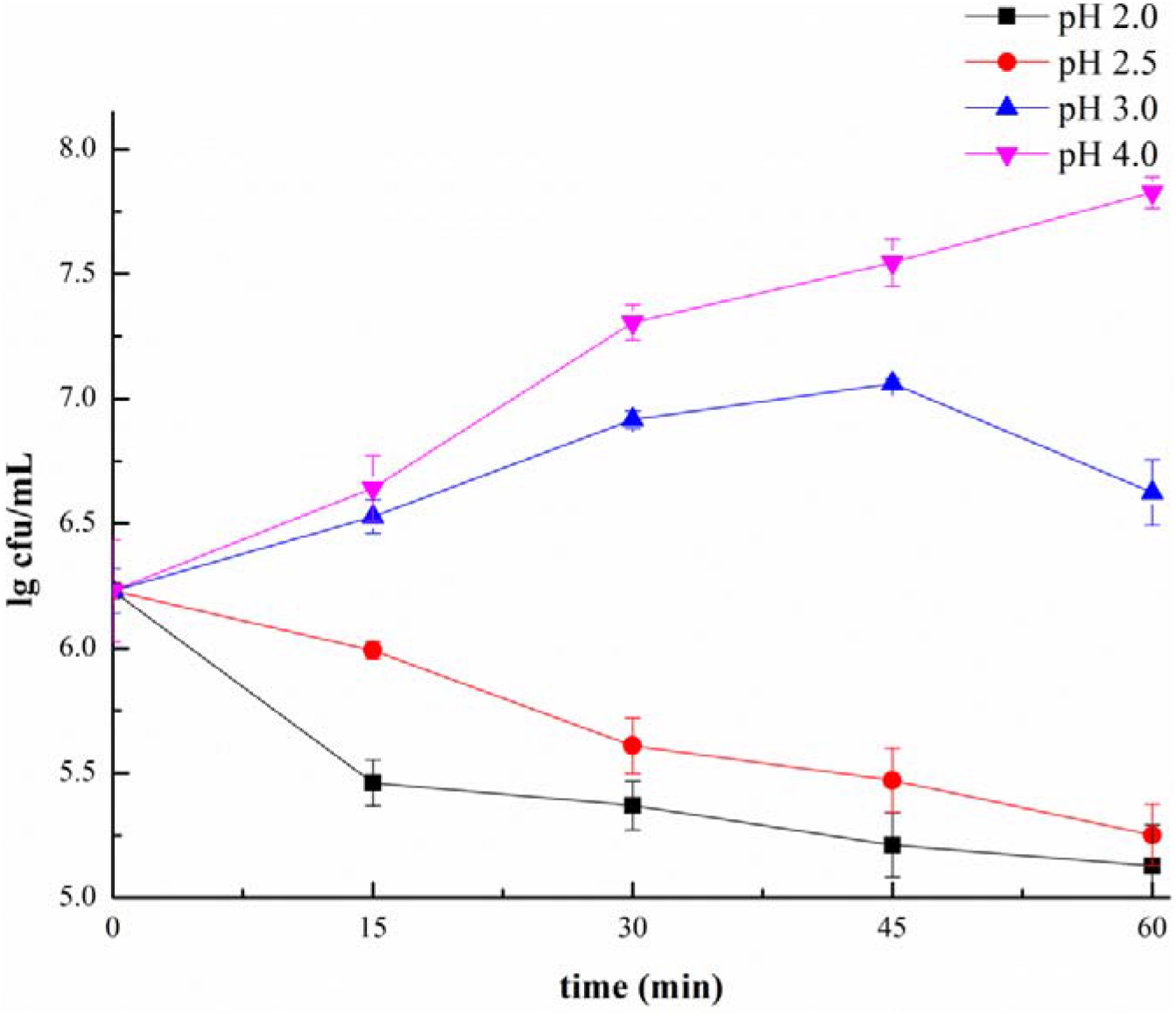
Survival curves of *A. acidoterrestris* under acid stress (different pH conditions: pH4.0, pH3, pH2.5, pH2.0, Under each pH, the sample exposed for 15-, 30-, 45- and 60-mins duration correspondingly. Data presentations were means ± SD from 3 replicates).

**Fig 2.**
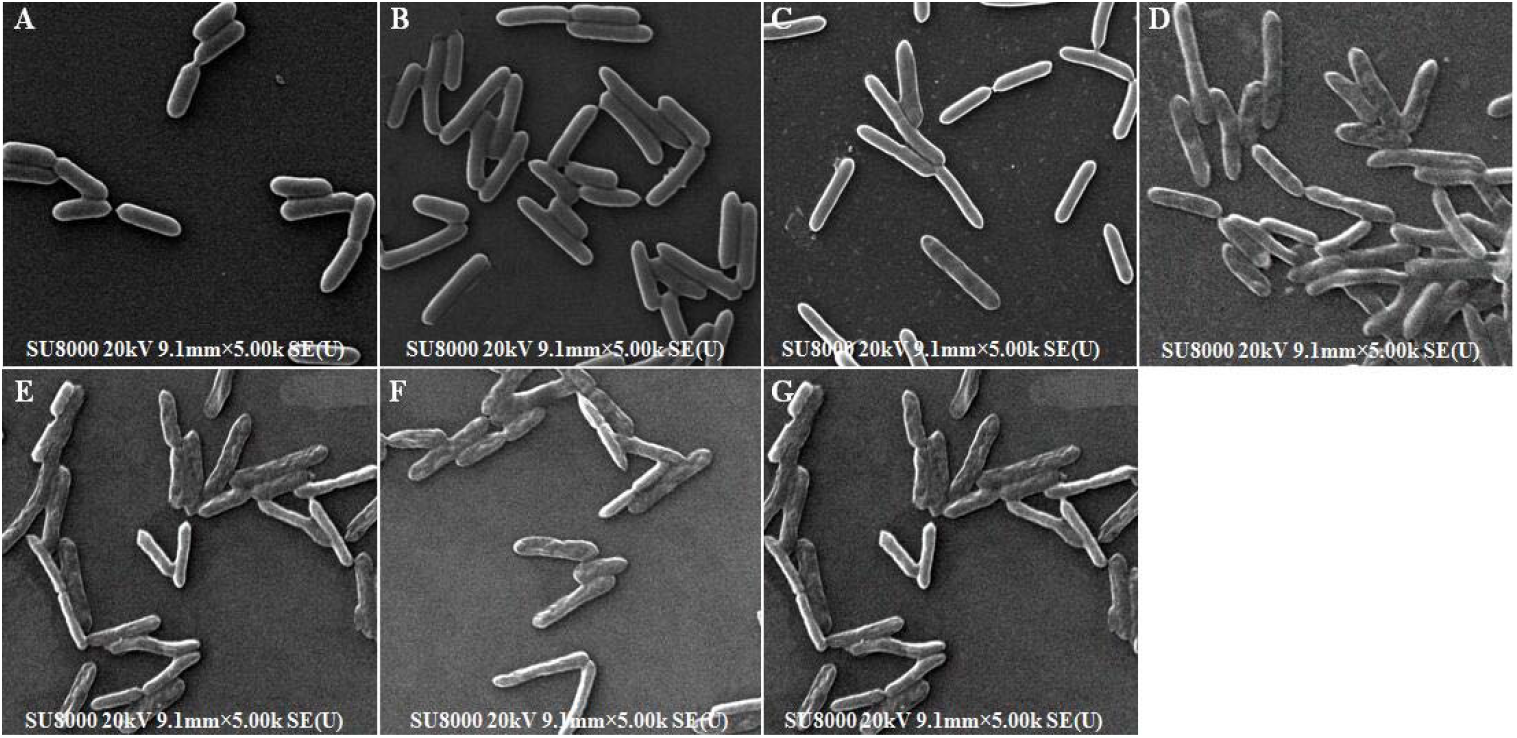
The SEM micrographs to show the morphology of *A. acidoterrestris* under different pH and time duration. A) Control, pH4.0, B-C) Acid stress at pH 2.0 for 15 min and 45 min, respectively, D-E) Acid stress at pH 2.5 for 15 min and 45 min, respectively, F-G) Acid stress at pH 3.0 for 15 min and 45 min, respectively.

### Identification of significantly changing of proteins expression level in *A. acidoterrestris* at acid stress

According to LC-ESI-MS/MS analysis and data processing results, significant analysis was performed for proteins that passed the *P*-value cutoff (P<0.05). In total, expression abundance of 325 proteins were significantly changed at acid stress for 15 min, among which 83 were observed to up-regulate their expression and 242 down-regulate their expression after pH2.5 acid stress for 15 min.

### Gene Ontology (GO) analysis

The differentially expressed proteins were analyzed using Gene Ontology (Fig. 3). The results showed that differential expressed proteins were involved in different biological process. The differentially expressed proteins mainly involved in metabolic process, cellular process, single-organism process, developmental process, cellular component organization or biogenesis, stimulus response, locomotion, negative regulation of biological process, reproduction, signalling and so on. Their molecular functions were involved in catalytic activity, structural molecule activity, transcription factor activity, metallochaperone activity, molecular function regulator, signal transducer activity, molecular transducer activity, electron carrier activity, nucleic acid binding transcription factor activity and transporter activity. In cellular component, those involved nine GO terms are cell, cell part, membrane, macromolecular complex, membrane part, organelle, organelle part, nucleoid, extracellular region. Among the differential expressed proteins, 237 were cell part, 85 were membrane and 51 were macromolecular complex and 25 were organelle.

**Fig 3.**
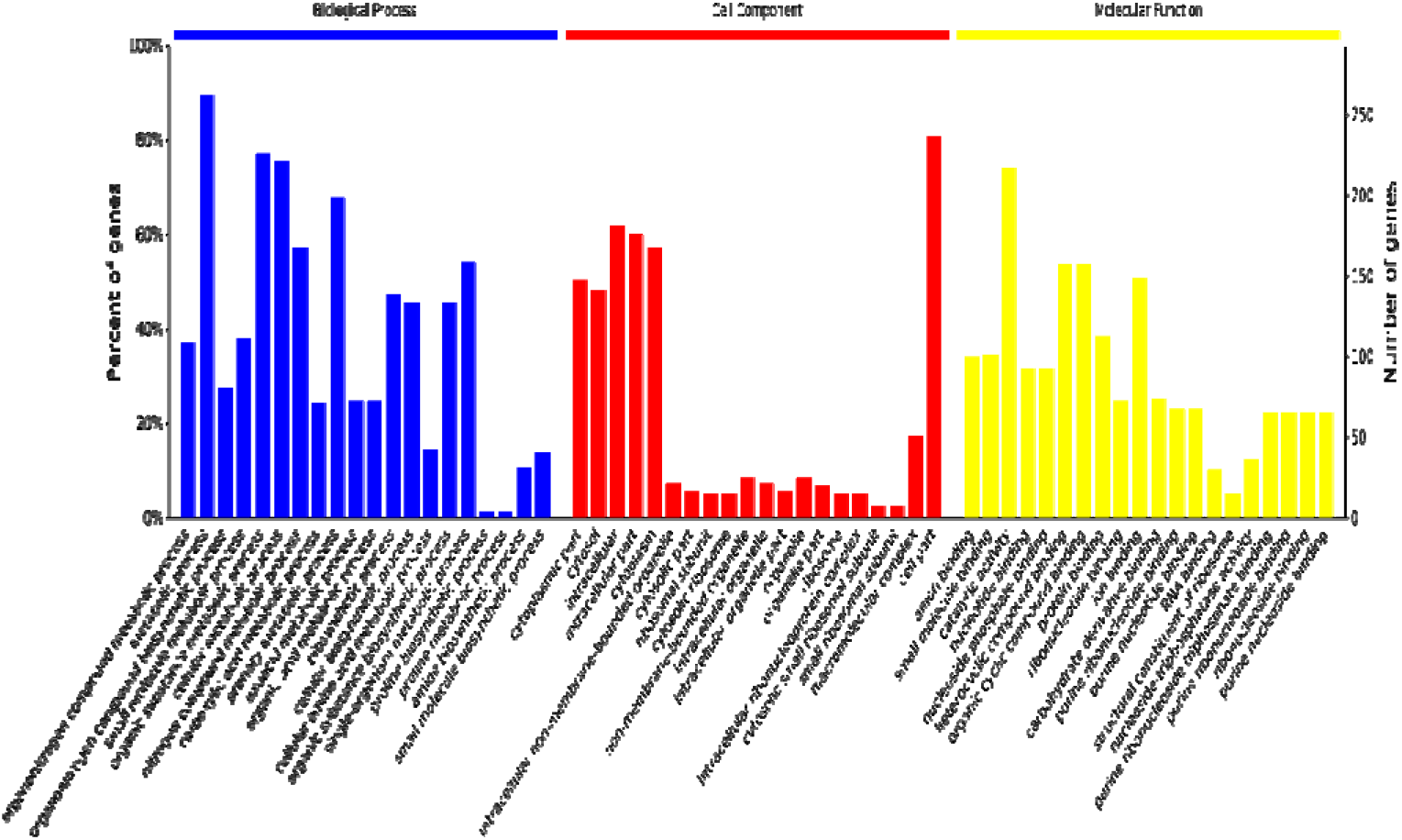
Analysis of differentially expressed proteins of *A. acidoterrestris* under acid stress. The differentially expressed proteins can mainly group into 3 classes: biological process, molecular function and cellular component.

### KEGG pathway analysis

Ninety-seven Kyoto Encyclopedia of Genes and Genomes pathways involving differential proteins were obtained through significant enrichment of pathway (*p*<0.05). Fig. 4 is the diagram of KEGG enrichment analysis of differential expressed proteins. Compared with the control group, biosynthesis of secondary metabolites, metabolic pathways and the carbon metabolism pathways related to the global and overview pathways, biosynthesis of unsaturated fatty acids, biosynthesis of antibiotics, biosynthesis of amino acids, mismatch repair and the pathways related to carbohydrate metabolism including glycolysis/gluconeogenesis, pentose phosphate pathway, citrate cycle were significantly enriched after acid stress. There were also some pathways, such as the two-component system, ABC transporters and bacterial secretion system, although there was no significant enrichment, there were still many up-regulated differential protein expressions.

**Fig 4.**
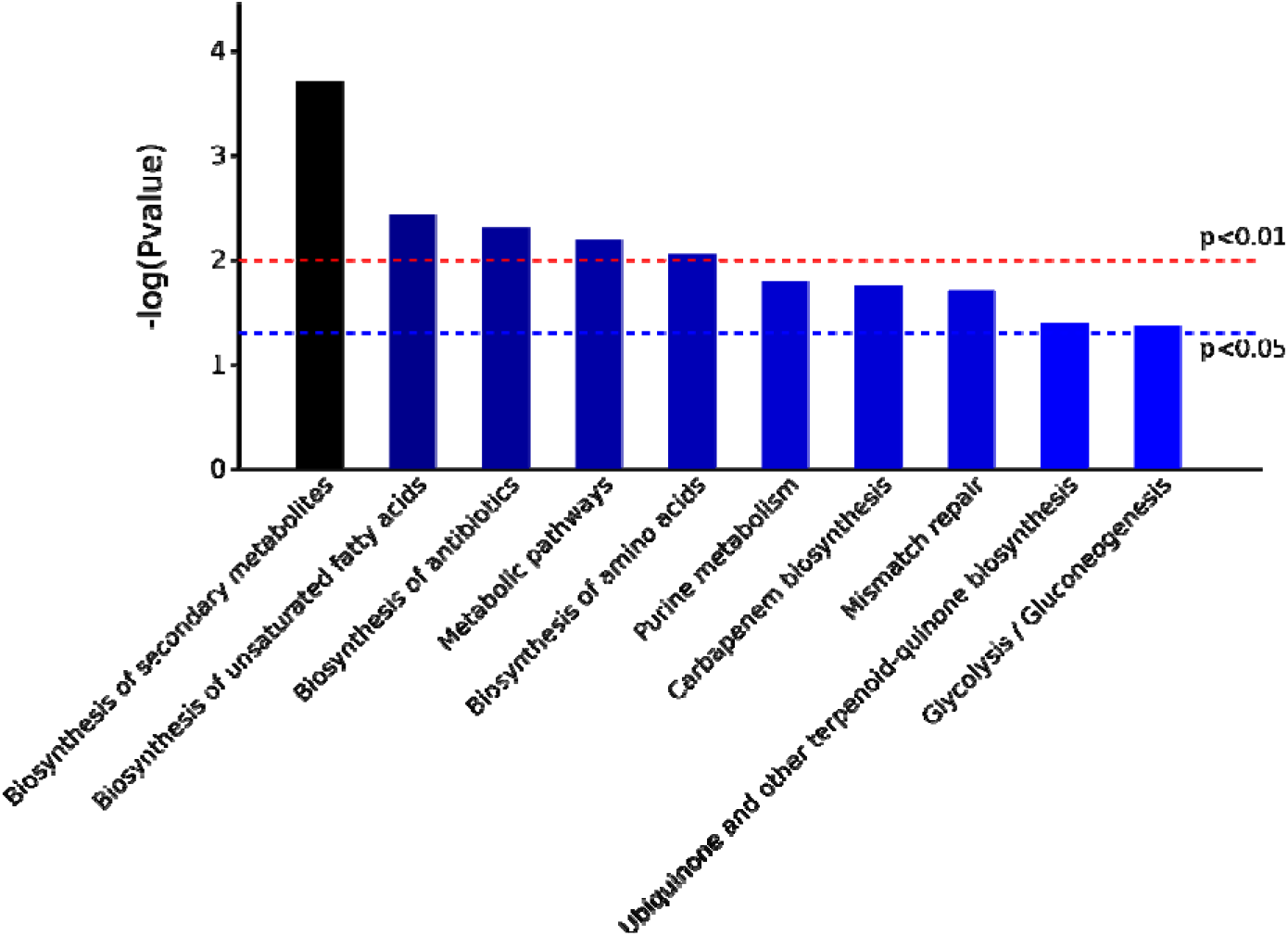
Top ten enriched pathways of the differentially expressed proteins of *A. acidoterrestris* under acid stress using KEGG pathway analysis.

**Fig 5.**
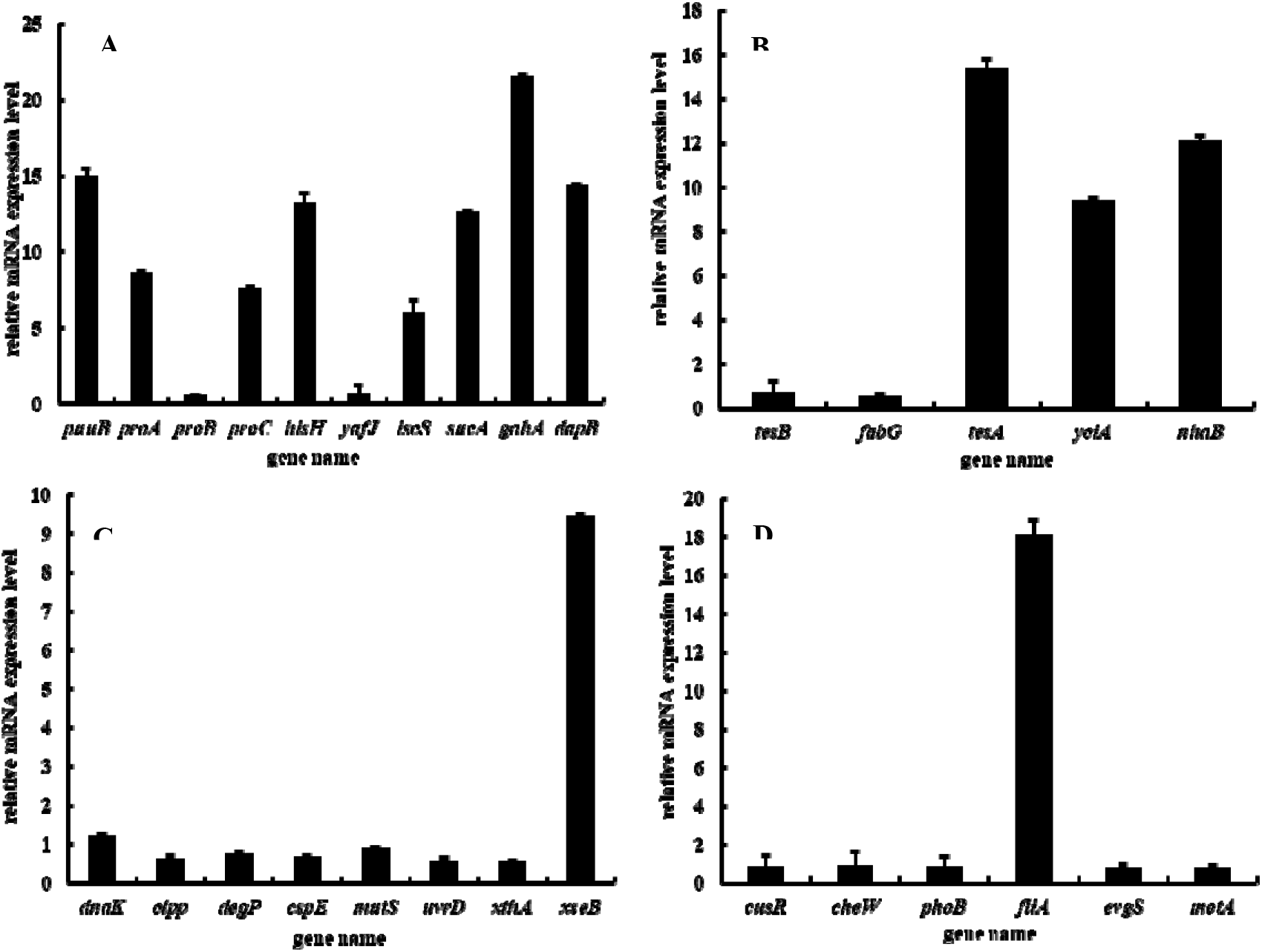
Relative mRNA expression levels of the significantly changed proteins. A). Proteins related to amino acid metabolism B). Proteins related to biosynthesis of unsaturated fatty acids C). Proteins related to stress response proteins and DNA damage repair D). Proteins related to two-component system (The results represent the mean ± SD of three biological replicates).

### Validation of differentially expressed proteins at mRNA level

Based on LFQ proteomic analysis results, a set of 29 significantly changed genes related to acid stress response were selected to be further validated using quantitative RT-PCR. The results of qRT-PCR were consistent with the protein expression trend of RNA seq, which verified the reliability of proteomic results. Among them, *proA, proC, iscS, sucA, dapB, hisH, gltD, gdhA* related to arginine, proline, lysine, histidine and glutamate metabolism were up-regulated. Except for *xseB*, other genes related to DNA repair including *mutS, uvrD, xthA*, and *endonuclease IV* were downregulated. Among the genes related to two-component regulatory system, all other genes including cusR, phoB and evgS are down-regulated, but *fliA* is up-regulated. The stress response protein including dnaK, clpP, and degP were down-regulated. The results showed that acid tolerance of *A. acidoterrestris* is regulated by multiple genes and different pathways.

## Discussion

The acid stress survival strategies of bacteria that involved in pH homeostasis mechanism including proton pumps, arginine decarboxylase (ADC) system, glutamate decarboxylase (GAD) system, lysine decarboxylase (LDC) system, and protection or repair of macromolecules such as proteins and DNA, the changes of cell membrane composition, cell density and biofilms, regulators, alteration of metabolic pathways and so on (Alvarez-Ordonez, Fernandez, Bernardo, & Lopez, 2010; Cotter & Hill, 2003; Krulwich, Sachs, & Padan, 2011). It is reported that the two most common ways bacteria generate alkali (ammonia, NH_3_) were involved in arginine deiminase (ADI) pathways and the urease. The production of NH_3_ combined with protons in the cytoplasm to produce NH_4_^+^, which raised the internal pH and increased the acid tolerance of cells (Cotter & Hill, 2003; Lindgren et al., 2014).

In the current study, the interesting discovery was that the expression level of NhaB was rapidly increased after 15 min of acid stress. NhaB and NhaA are two major Na^+^/H^+^ antiporters found in *Escherichia coli*, which can sense and transmit external pH signals and play a crucial role in maintaining intracellular pH homeostasis. NhaA is only activated when the intracellular pH reaches 6.5 (Padan & Dwivedi, 2015), while NhaB mainly functions in lower pH environments, collaborating with NhaA proteins to participate in environmental stress responses(Pinner, Kotler, & Padan, 1993). As a kind of obligate acidophilic bacteria, it is unclear how *A. acidoterrestris* perceive the external pH environment. The results indicated NhaB protein was closely related to the perception of extracellular pH levels by *A. acidoterrestris*.

Moreover, 5 differential proteins were mapped into arginine and proline metabolism pathway under acid stress, among which arginase, γ-glutamyl phosphate reductase (proA) and pyrroline-5-carboxylate reductase (proC) were found significantly up-regulated. Arginine can be degraded by three enzymes: arginase, arginine decarboxylase (ADC) and arginine deiminase (ADI) (Hsueh et al., 2012). It is well-known that arginase converts L-arginine to L-ornithine and urea, and catalyzes the fifth and final step in the urea cycle and is part of a pathway for detoxifying ammonia (Wu & Morris, 1998). However, the increased expression of arginase reported here indicated that it is induced by the reduction of pH and the promotion of the ammonia production in theory was reminiscent of its responsibility for acid-tolerance in *A. acidoterrestris*. There exists two isozymes of arginase, arginase I functions in the urea cycle and arginase II regulates the arginine/ornithine concentrations in the cell (Morris, 2002). And then, arginine/ornithine concentrations modulated the reverse transport of arginine-ornithine (Tonon & Lonvaud-Funel, 2000). Although a previous study proposed that ADI pathway is far more important than urease for acid resistance of in *Laribacter hongkongensis* (Xiong et al., 2014), it is possible that *A. acidoterrestris* sharing inherent acid resistance may contain a special arginine metabolic system in respond to acid stress. Its role in acid resistance of *A. acidoterrestris* is largely unknown and is to be further investigated. It is rather remarkable that the increase of proA and proC related to synthesizing of L-proline from L-glutamate suggested that the synthesis of L-proline was raised at acid stress. L-glutamate is also a precursor of L-arginine synthesis and the initial step namely N-acetylation of L-glutamate in the synthesis of arginine prevents the spontaneous cyclization of the glutamate semialdehyde intermediate, thus precludes the formation of the proline precursor pyrroline-5-carboxylate and further synthesis of proline (Errey & Blanchard, 2005). So, we could infer that metabolic path of glutamine in *A. acidoterrestris* at acid stress were precisely regulated and glutamine mainly flowed into the arginine synthesis pathway based on the data reported here, which provided proteomics evidence for bacteria by alter amino acid metabolic pathways to adapt to low pH environments.

Amino acid-dependent acid resistance systems have been clearly defined in *E. coli*. The arginine decarboxylase (ADC) decarboxylates arginine to agmatine in a mechanism that is similar to that of glutamate decarboxylase and the two decarboxylase systems are believed to consume intracellular protons and release CO_2_ during the decarboxylation of glutamate or arginine, thus protect *E. coli* against acid stress (Castanie-Cornet et al., 2010; Castanié-Cornet, Penfound, Smith, Elliott, & Foster, 1999). Subsequently, the resistance to acid stress of two decarboxylase systems were described in a variety of bacteria(Fernandez & Zuniga, 2006; Lin, Lee, Frey, Slonczewski, & Foster, 1995; Valenzuela et al., 2014). Moreover, histidine decarboxylation system was identified in lactic acid bacteria to function like the two decarboxylates systems and improve acid stress survival of *Lactococcus lactis* (Broadbent, Larsen, Deibel, & Steele, 2010; Fernandez & Zuniga, 2006; Trip, Mulder, & Lolkema, 2012). It has been widely accepted that amino acid decarboxylation is another important way for bacteria to resist acid stress. However, the three AR system described for *E. coli* did not present in *Salmonella typhimurium* (Lin et al., 1995). It can be inferred that bacteria with different acid resistance also have evolved different acid-resistant mechanisms. In *A. acidoterrestris*, the decrease of the enzymes related to glutamate decarboxylation and the increase of the histidine metabolism-associated proteins after acid stress inclined that there might exist a similar histidine-dependent acid resistance system to lactic acid bacteria.

In recent years, many scholars have discovered and identified protective proteins related to pH pressure. Most of the identified acid stress proteins involved in cell regulation, energy metabolism, molecular monitoring, transcription, translation, pili synthesis, cell signaling, and virulence(Ritter et al., 2014). It is reported that bacteria may activate different acid resistant systems under different pH conditions. In *Escherichia coli*, its five acid resistant systems that could keep it survive for hours at around pH 2, but have no effect on the growth of *Escherichia coli* at pH 4-6 (Y. Xu et al., 2020). In this study, the expression levels of stress proteins, including clpP, dnaK, and DegP, were downregulated. It is speculated that the acid resistance system mediated by these stress proteins is not the main acid resistance mechanism of *A. acidoterrestris* at sublethal pH conditions.

The enzymes related to DNA mismatch repair and base excision repair such as mutS, uvrD, xseB and xthA changed dramatically in the study. DNA damage caused by acid has been confirmed in a variety of bacteria including *Escherichia coli, Bacillus subtilis, Streptococcus* and so on (Hanna, Ferguson, Li, & Cvitkovitch, 2001; Jeong, Hung, Baumler, Byrd, & Kaspar, 2008; Lindahl & Nyberg, 1972). It has shown that low pH values could result in DNA double-strand breaks of bacteria and lead to the depurination of the DNA, consisting in the loss of purines which results in the formation of apurinic sites (Jeong et al., 2008; Lindahl & Nyberg, 1972). Accordingly, loss of genetic information caused by the depurination induced DNA repair systems in bacteria. Indeed, several bacterial mutants of the DNA repair systems showed heightened acid sensitivity, consistent with DNA damage being a consequence of acidification (Hanna et al., 2001; Quivey, Faustoferri, Clancy, & Marquis, 1995). DNA repair systems are ubiquitous and essential to all living organisms. One of the most important DNA repair systems in bacteria is the Dam-dependent mismatch repair system (MMR) including MutS, MutH and MutL. MutS recognizes a mismatched base-pair as well as insertions or deletions of one to four nucleotides. MutL forms a complex with MutS that activates the MutH endonuclease, and the *uvrD* gene product known as DNA helicase II ensures the required separation of DNA strands (Matic, Rayssiguier, & Radman, 1995; Rayssiguier, Thaler, & Radman, 1989). After acid stress at pH2.5 for 15min, the expression of MutS, UvrD and exodeoxyribonuclease III encoded by *xthA* was down-regulated dramatically, and *xseB* was up-regulated notably. The changes were validated from transcriptional level by qRT-PCR. The increase of *xseB* indicated that *xseB* involved in DNA repair by nucleotide excision which might played a crucial role in repairing acid-induced DNA damage in *A. acidoterrestris* under acid stress. Excision repair relies on the redundant information in the duplex to remove a damaged base or nucleotide and replace it with a normal base by using the complementary strand as a template (Sancar, 1994). It has been suggested that the role of base excision repair involving the AP endonuclease that could repair minor DNA damage whereas UvrA family and the nucleotide excision repair pathway could be responsible for excising larger DNA lesions caused by acid and other DNA-damaging agents (Cotter & Hill, 2003). Although which types of DNA damage caused by acid stress at pH 2.5 in *A. acidoterrestris* remains unclear, the significant changes of these enzymes indicated that the DNA damage types might affect the specific expression of enzymes.

Cell membrane is one of the primary targets of physical and chemical factors from the external environment on which bacteria rely for survival. Bacteria can protect cells by changing the structure, composition, stability, and activity of cell membranes. The changes in the fatty acid composition of cell membranes are the most common response of bacteria to external environmental pressure. The newly discovered acid tolerance mechanism of *Escherichia coli* indicated that it could directly sense the acidic environment through the sensing kinase CpxA and initiated the expression of unsaturated fatty acid synthesis genes, thereby increasing the proportion of unsaturated fatty acids in cell membrane phospholipids, reducing the fluidity and cytoplasmic permeability of cell membranes, and maintaining the pH inside *Escherichia coli* cells free from the influence of external acidic environments (Cotter & Hill, 2003; Y. Xu et al., 2020). Under acid stress, the pathway of biosynthesis of unsaturated fatty acids in *A. acidoterrestris* was significantly enriched, with the expression levels of tesA protein and yciA protein upregulated by 1.54 and 1.81 times, respectively, indicating that regulation of cell membrane permeability is also the main means for *A. acidoterrestris* to respond to extremely low pH environments.

The results of the study showed that *A. acidoterrestris* rapidly adjusted its protein expression levels, metabolic pathways and membrane permeability to adapt to acid stress environments at pH 2.5. We know that bacterial response to external acid stress environments is a complex mechanism. How to sense and transmit extracellular pH signals and how to coordinate these acid stress response proteins and metabolic pathways for maintaining intracellular pH homeostasis remains uncovered. Continuing exploration of their internal relationships and regulatory mechanisms will provide new genetic materials and means for strain resistance cultivation in the fermentation industry.

## Acknowledgements

This work was supported by National Natural Science Foundation of China (Grant numbers 31771949, awarded to LJ) and Program for Science and Technology Innovation Talents in Universities of Henan Province (Grant numbers 17HASTIT037, awarded to LJ).

## Ethical Statement

This article does not contain any studies with human participants or animals performed by any of the authors.

## Conflict of Interest

All the authors declare that they have no conflict of interest.

## Notes

### Competing Interest Statement

The authors have declared no competing interest.

